# Octapeptin C4 Induces Less Resistance and Novel Mutations in an Epidemic Carbapenemase-producing *Klebsiella pneumoniae* ST258 Clinical Isolate Compared to Polymyxins

**DOI:** 10.1101/309674

**Authors:** Miranda E. Pitt, Minh Duc Cao, Mark S. Butler, Soumya Ramu, Devika Ganesamoorthy, Mark A. T. Blaskovich, Lachlan J. M. Coin, Matthew A. Cooper

## Abstract

Polymyxin B and E (colistin) have been pivotal in the treatment of extensively drug-resistant (XDR) Gram-negative bacterial infections, with increasing use over the past decade. Unfortunately, resistance to these antibiotics is rapidly emerging. The structurally-related octapeptin C4 (OctC4) has shown significant potency against XDR bacteria, including against polymyxin-resistant (Pmx-R) strains, but its mode of action remains undefined. We sought to compare and contrast the acquisition of XDR *Klebsiella pneumoniae* (ST258) resistance *in vitro* with all three lipopeptides to help elucidate the mode of action of the drugs and potential mechanisms of resistance evolution. Strikingly, 20 days of exposure to the polymyxins resulted in a dramatic (1000-fold) increase in the minimum inhibitory concentration (MIC) for the polymyxins, reflecting the evolution of resistance seen in clinical isolates, whereas for OctC4 only a 4-fold increase was witnessed. There was no cross-resistance observed between the polymyxin - and octapeptin-induced resistant strains. Sequencing revealed previously known gene alterations for polymyxin resistance, including *crrB*, *mgrB*, *pmrB*, *phoPQ* and *yciM*, and novel mutations in *qseC*. In contrast, mutations in *mlaDF* and *pqiB*, 1genes related to phospholipid transport, were found in octapeptin-resistant isolates. Mutation effects were validated via complementation assays. These genetic variations were reflected in phenotypic changes to lipid A. Pmx-R isolates increased 4-amino-4-deoxy-arabinose fortification to phosphate groups of lipid A, whereas OctC4 induced strains harbored a higher abundance of hydroxymyristate and palmitoylate. The results reveal a differing mode of action compared to polymyxins which provides hope for future therapeutics to combat the increasingly threat of XDR bacteria.

## INTRODUCTION

Infections by extensively drug-resistant (XDR) bacteria are an increasing concern due to the lack of effective antibiotics, thereby resulting in high mortality (1, 2). Common therapeutic interventions include fosfomycin, tigecycline and polymyxins (2-4). However, the effectiveness of these therapies is short lived due to plasmid-encoded resistance (fosfomycin (4), polymyxin (5)) and rapid acquisition of resistance through mutation (fosfomycin (6), tigecycline (7) and polymyxin (8, 9)). New antibiotics with the capacity to ablate these XDR bacteria are urgently desired.

Octapeptins are structurally similar to the polymyxins, with both lipopeptide classes consisting of a cyclic heptapeptide ring and linear tail capped with a fatty acid, containing multiple positively charged diaminobutyric acid (Dab) residues (10-12) (Fig. 1). Studies on the polymyxins have shown that these Dab residues are critical for interactions with the basal component of lipopolysaccharide (LPS), lipid A. The mode of action involves the initial binding to lipid A, displacement of magnesium (Mg^2+^) and calcium (Ca^2+^), permeabilization of the outer and inner membrane, leakage of cytoplasmic contents and subsequent cell death, however, the exact mechanism is yet to be discerned (13, 14). The phosphate groups on lipid A are modified during polymyxin resistance with 4-amino-4-deoxy-arabinose (Ara4N) and/ or phosphoethanolamine (pEtN) in order to stabilise the outer membrane. This reduces polymyxin binding by removing the negative phosphate that attracts the cationic Dab residues (15, 16). Constitutive up-regulation of this pathway is achieved through chromosomal variations in the two-component regulatory systems (TCS) *crrAB, pmrAB, phoPQ* and the negative regulator *mgrB* in *Klebsiella pneumoniae* (8, 9, 17). These modifications perturb the electrostatic interaction between lipid A and polymyxins to negate the infiltration of this antibiotic class. The structurally similar octapeptins retain most of the key binding motifs, and might be expected to employ a similar mode of action. The most significant structural difference between the polymyxins and octapeptins is a truncated linear exocyclic peptide (1 residue instead of three) linked to a β-hydroxy-fatty acid (instead of an alkyl fatty acid) in the octapeptins (10-12). More minor variations include L-Dab to D-Dab and L-Thr to L-Leu substitutions. Loss of the fatty acid tail component has been shown to attenuate activity in polymyxins (18). Despite their similarity, prior research has revealed octapeptins retain the ability to kill Pmx-R bacteria, inferring an alternative mode of action (10). In addition, some octapeptins have broad spectrum activity with potency against Gram-positive bacteria, fungi and protozoa (19, 20).

**FIG.1.**
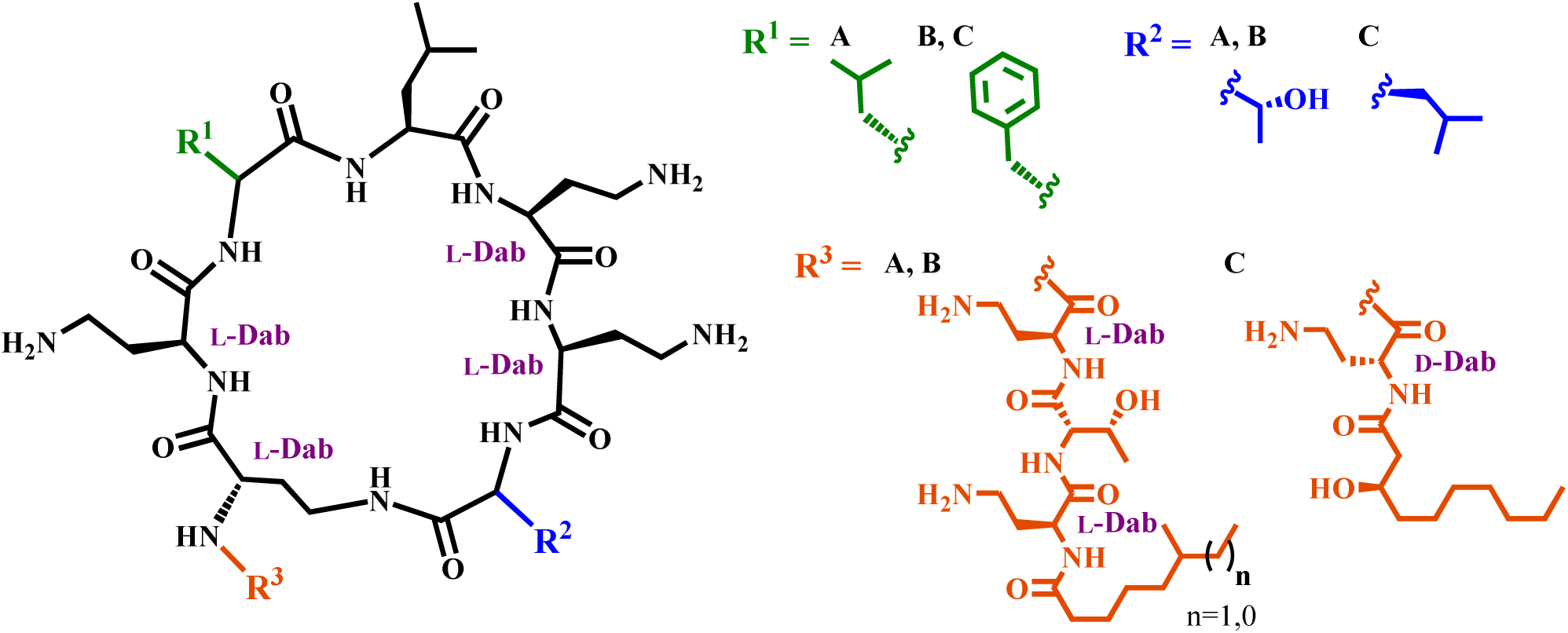
Structural comparison between the 3 lipopeptide antibiotics used in this study. Colistin (A) and polymyxin B (B) differ by one amino acid (polymyxin B: phenylalanine and colistin: leucine (R^1^)). One defining feature of octapeptin C4 (C) is that it contains 8 amino acids rather than 10 in polymyxins. In addition, a leucine residue replaces threonine within the ring (R^2^), the exocyclic diaminobutyric acid (Dab) residue is the D-enantiomer, and the fatty acid tail contains a 3-hydroxy group (R^3^).

We have recently reported the first synthesis of Octapeptin C4 (OctC4) (10) and Octapeptin A3 (21)followed by detailed biological characterisation of OctC4 that demonstrates its potential as a new ‘last resort’ antibiotic to treat serious extensively drug-resistant Gram-negative infections. (22)In view of the limited understanding of the mechanism by which octapeptins target bacteria, we sort to investigate the differences driving development of OctC4 and polymyxin resistance at a genetic level. Two studies have previously investigated the acquisition of resistance towards octapeptins. One was performed using EM49 (a mixture of octapeptin classes A and B) which exhibited no increase in resistance after 10 passages for *Pseudomonas aeruginosa, Escherichia coli, Staphylococcus aureus* and *Candida albicans* (23). The other investigated lipid A modifications in *P. aeruginosa* isolates resistant to OctC4 obtained from a subculture surviving a single overnight treatment at 2 or 32 μg/ml.

The ST258 lineage of *K. pneumoniae* is endemic in numerous regions in the world and commonly involved in outbreaks (24-27). These isolates pose as a major threat due to frequently harboring carbapenem resistance, predominantly facilitated via *blaKPC* genes encoded on plasmids (25, 28). We have previously used whole genome sequencing to investigate the acquisition of resistance to polymyxin in an endemic lineage of *K. pneumoniae*, ST258, isolated from a Greek hospital (24). We selected one of these isolates which is susceptible to polymyxin and OctC4, but otherwise highly resistant (*aac(6’)Ib, aac(6’)Ib-cr, aph(3’)-Ia, aph(3”)-Ib, aph(6)-Id, blaKPC-2, blaLEN-*12* blaOXA-9, blaTEM-1B, fosA, oqxAB, sul2, tet(A), dfrA14* positive), for resistance induction experiments. This strain is representative of the type of pathogens for which a ‘last resort’ antibiotic is employed, and one where high levels of clinical resistance to polymyxins have already been well characterised at a genetic level (24). The resistance induction experiments were followed by characterization of antibiotic susceptibility, whole genome sequencing, and analysis of lipid A composition. This research has uncovered significant differences in the development of resistance induced by the two classes of antibiotics, both in the level of resistance created and in the underlying genetic mutations. The results provide strong support for further development of the octapeptins as potential last-resort therapeutics.

## RESULTS

### Rapid resistance acquisition for polymyxins dissimilar to OctC4

Forced evolution on XDR *K. pneumoniae* was monitored over a 20 day time course, with six replicates exposed to increasing concentrations of either polymyxin B (PMB), polymyxin E (colistin, CST) and OctC4. Significant variability was observed between polymyxins and OctC4 (Fig. 2D). Initially, an MIC of 0.125 μg/ml was measured for both CST and PMB. The majority of replicates treated with the polymyxins had a clinical resistance phenotype of >2 μg/ml by day 10 (Fig. 2A and B). At some point, every replicate had a dramatic and rapid escalation in MIC to >64 μg/ml, generally over ≤5 days. The timing for the drastic increase varied between replicates, and appeared dependent on whether the replicate could tolerate 0.5 μg/ml, with a steep escalation from the day that resistance level was exceeded. In sharp contrast, OctC4 resistance progressed steadily over the 20 days (Fig. 2C) with only a 4-fold (initial MIC: 8 μg/ml) overall increase compared to a ≥1000-fold increase for the polymyxins (Fig. 2E). The trend for a gradual increase in OctC4 resistance was consistent amongst replicates. The induced resistance appeared to be stable after five additional passages without antibiotic exposure for polymyxins. The extent of growth in wells containing either 32 or 16 μg/ml started to diminish for OctC4-induced isolates during the last passages.

**FIG.2.**
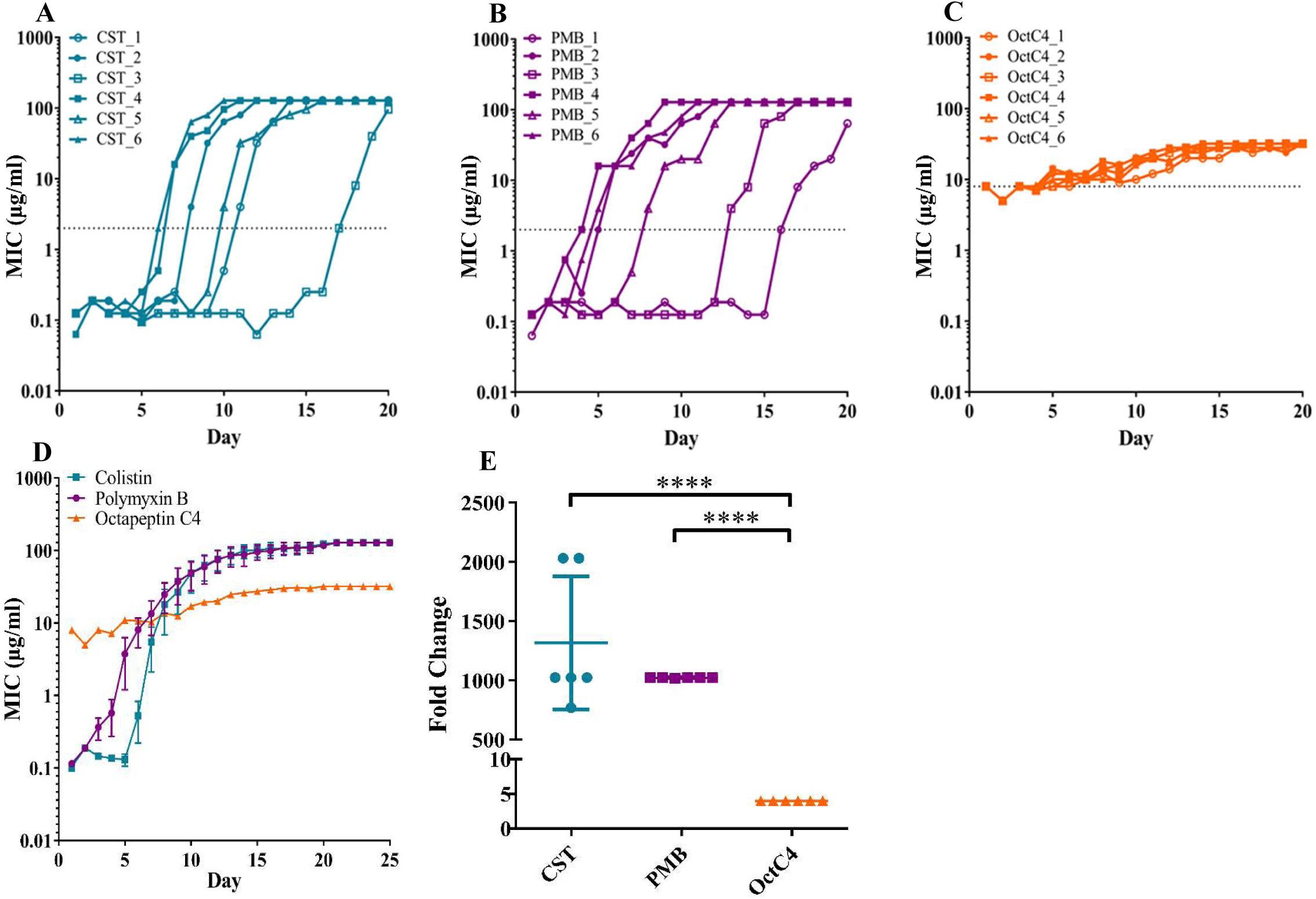
Acquired resistance in extensively drug-resistant *K. pneumoniae* over time for polymyxins and octapeptin C4. (A) Colistin (B) Polymyxin B (C) Octapeptin C4 (D) Overall comparison of acquired resistance for 20 day antibiotic exposure and 5 days following without exposure (mean±SEM, n=6) (E) Fold change of colistin (CST), polymyxin B (PMB), and octapeptin C4 (0ctC4) in concordance to day 0 and 20 MIC (mean±SD) (^* * * *^P<0.001). Line represents break points (2 μg/ml polymyxins, 8 μg/ml set for octapeptin to highlight divergence from day 0). Highest concentration used for polymyxins was 128 μg/ml and 32 μg/ml for octapeptin C4.

### Lack of cross-reactivity and reduction of resistance in OctC4 induced isolates

Day 20 isolate MICs were determined against a broad array of antibiotic classes to evaluate if acquired resistance conferred cross-resistance, or resulted in regained susceptibility (Table 1). Remarkably, no cross-reactivity was apparent between polymyxins and OctC4. Non-susceptibility towards amoxicillin, aztreonam, ceftriaxone, ciprofloxacin, piperacillin and trimethoprim was ubiquitous amongst treatment groups. Chloramphenicol resistance was observed in the initial isolate but was diminished in the majority of replicates over the time course for all three antibiotics. In some instances, cefepime susceptibility was restored (replicates OctC4_3, OctC4_4 and OctC4_6). These replicates also regained susceptibility to meropenem, as did PMB_2. Replicate OctC4_2 also exhibited susceptibility to tetracycline and tigecycline whilst this profile for OctC4_4 varied extensively for these antibiotics, where resistant and susceptible MICs were recorded depending on the colonies selected.

**TABLE 1.**
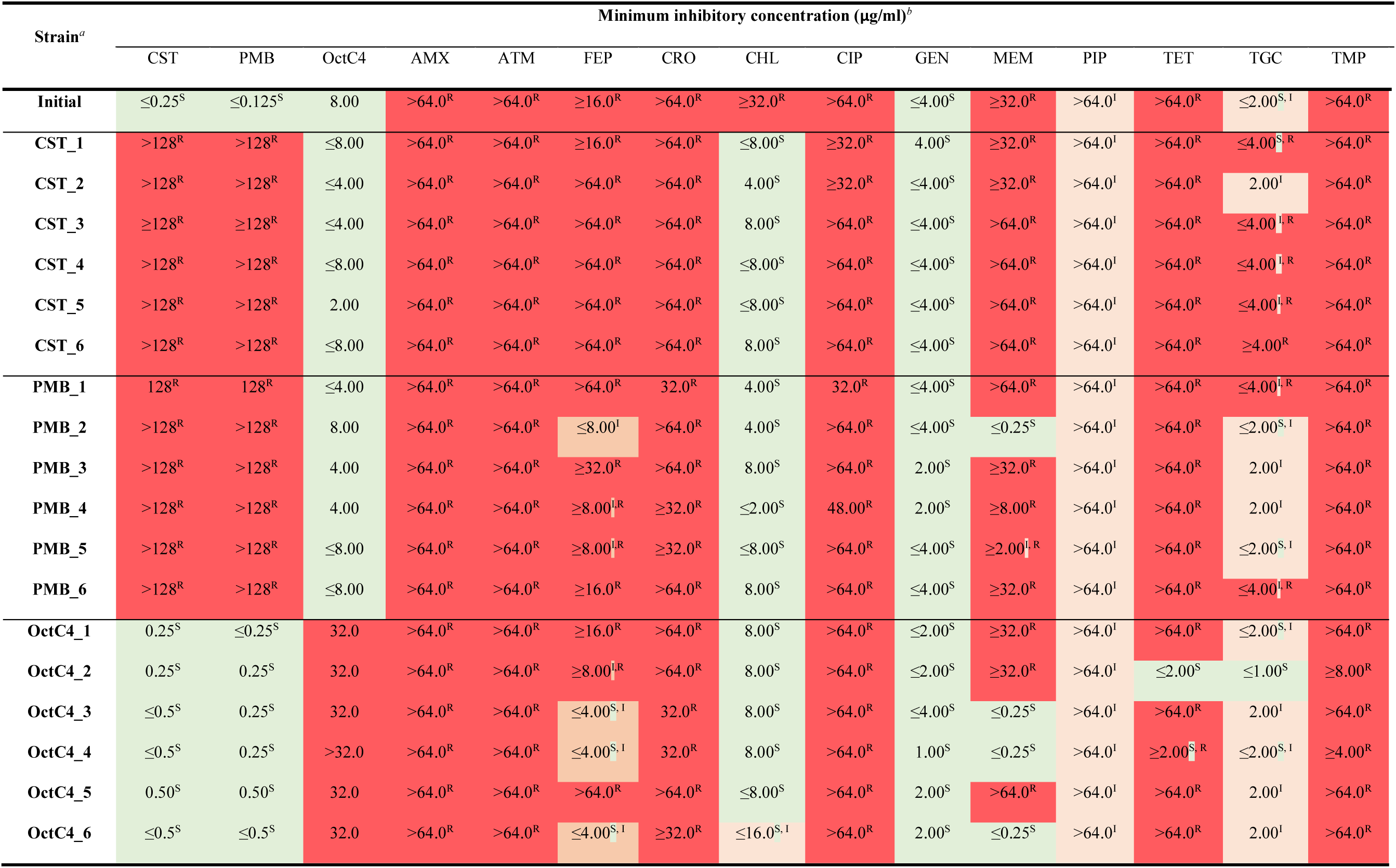
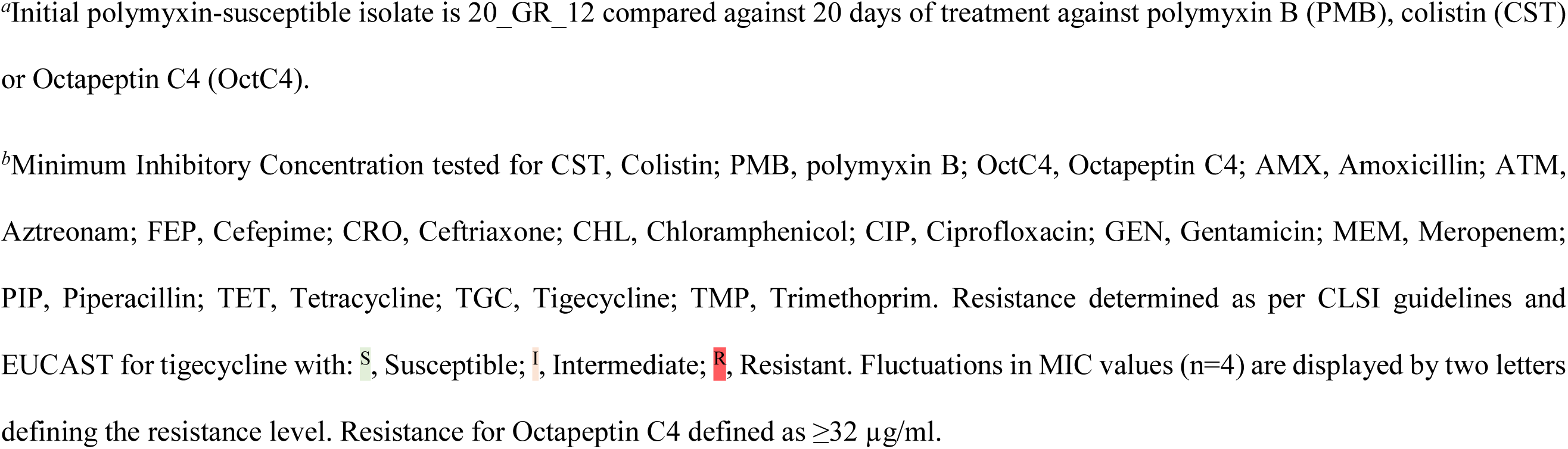
Minimum inhibitory concentrations of day 20 replicates compared to initial isolate across several antibiotic classes

| Strain<sup>a</sup> | Minimum inhibitory concentration (µg/ml)<sup>b</sup> |  |  |  |  |  |  |  |  |  |  |  |  |  |  |
| --- | --- | --- | --- | --- | --- | --- | --- | --- | --- | --- | --- | --- | --- | --- | --- |
|  | CST | PMB | OctC4 | AMX | ATM | FEP | CRO | CHL | CIP | GEN | MEM | PIP | TET | TGC | TMP |
| **Initial** | ≤0.25<sup>S</sup> | ≤0.125<sup>S</sup> | 8.00 | >64.0<sup>R</sup> | >64.0<sup>R</sup> | ≥16.0<sup>R</sup> | >64.0<sup>R</sup> | ≥32.0<sup>R</sup> | >64.0<sup>R</sup> | ≤4.00<sup>S</sup> | ≥32.0<sup>R</sup> | >64.0<sup>I</sup> | >64.0<sup>R</sup> | ≤2.00<sup>S,I</sup> | >64.0<sup>R</sup> |
| **CST_1** | >128<sup>R</sup> | >128<sup>R</sup> | ≤8.00 | >64.0<sup>R</sup> | >64.0<sup>R</sup> | ≥16.0<sup>R</sup> | >64.0<sup>R</sup> | ≤8.00<sup>S</sup> | ≥32.0<sup>R</sup> | 4.00<sup>S</sup> | ≥32.0<sup>R</sup> | >64.0<sup>I</sup> | >64.0<sup>R</sup> | ≤4.00<sup>S,R</sup> | >64.0<sup>R</sup> |
| **CST_2** | >128<sup>R</sup> | >128<sup>R</sup> | ≤4.00 | >64.0<sup>R</sup> | >64.0<sup>R</sup> | >64.0<sup>R</sup> | >64.0<sup>R</sup> | 4.00<sup>S</sup> | ≥32.0<sup>R</sup> | ≤4.00<sup>S</sup> | ≥32.0<sup>R</sup> | >64.0<sup>I</sup> | >64.0<sup>R</sup> | 2.00<sup>I</sup> | >64.0<sup>R</sup> |
| **CST_3** | ≥128<sup>R</sup> | ≥128<sup>R</sup> | ≤4.00 | >64.0<sup>R</sup> | >64.0<sup>R</sup> | >64.0<sup>R</sup> | >64.0<sup>R</sup> | 8.00<sup>S</sup> | >64.0<sup>R</sup> | ≤4.00<sup>S</sup> | >64.0<sup>R</sup> | >64.0<sup>I</sup> | >64.0<sup>R</sup> | ≤4.00<sup>I,R</sup> | >64.0<sup>R</sup> |
| **CST_4** | >128<sup>R</sup> | >128<sup>R</sup> | ≤8.00 | >64.0<sup>R</sup> | >64.0<sup>R</sup> | >64.0<sup>R</sup> | >64.0<sup>R</sup> | ≤8.00<sup>S</sup> | >64.0<sup>R</sup> | ≤4.00<sup>S</sup> | >64.0<sup>R</sup> | >64.0<sup>I</sup> | >64.0<sup>R</sup> | ≤4.00<sup>I,R</sup> | >64.0<sup>R</sup> |
| **CST_5** | >128<sup>R</sup> | >128<sup>R</sup> | 2.00 | >64.0<sup>R</sup> | >64.0<sup>R</sup> | >64.0<sup>R</sup> | >64.0<sup>R</sup> | ≤8.00<sup>S</sup> | >64.0<sup>R</sup> | ≤4.00<sup>S</sup> | >64.0<sup>R</sup> | >64.0<sup>I</sup> | >64.0<sup>R</sup> | ≤4.00<sup>I,R</sup> | >64.0<sup>R</sup> |
| **CST_6** | >128<sup>R</sup> | >128<sup>R</sup> | ≤8.00 | >64.0<sup>R</sup> | >64.0<sup>R</sup> | >64.0<sup>R</sup> | >64.0<sup>R</sup> | 8.00<sup>S</sup> | >64.0<sup>R</sup> | ≤4.00<sup>S</sup> | >64.0<sup>R</sup> | >64.0<sup>I</sup> | >64.0<sup>R</sup> | ≥4.00<sup>R</sup> | >64.0<sup>R</sup> |
| **PMB_1** | 128<sup>R</sup> | 128<sup>R</sup> | ≤4.00 | >64.0<sup>R</sup> | >64.0<sup>R</sup> | >64.0<sup>R</sup> | 32.0<sup>R</sup> | 4.00<sup>S</sup> | 32.0<sup>R</sup> | ≤4.00<sup>S</sup> | >64.0<sup>R</sup> | >64.0<sup>I</sup> | >64.0<sup>R</sup> | ≤4.00<sup>I,R</sup> | >64.0<sup>R</sup> |
| **PMB_2** | >128<sup>R</sup> | >128<sup>R</sup> | 8.00 | >64.0<sup>R</sup> | >64.0<sup>R</sup> | ≤8.00<sup>I</sup> | >64.0<sup>R</sup> | 4.00<sup>S</sup> | >64.0<sup>R</sup> | ≤4.00<sup>S</sup> | ≤0.25<sup>S</sup> | >64.0<sup>I</sup> | >64.0<sup>R</sup> | ≤2.00<sup>S,I</sup> | >64.0<sup>R</sup> |
| **PMB_3** | >128<sup>R</sup> | >128<sup>R</sup> | 4.00 | >64.0<sup>R</sup> | >64.0<sup>R</sup> | ≥32.0<sup>R</sup> | >64.0<sup>R</sup> | 8.00<sup>S</sup> | >64.0<sup>R</sup> | 2.00<sup>S</sup> | ≥32.0<sup>R</sup> | >64.0<sup>I</sup> | >64.0<sup>R</sup> | 2.00<sup>I</sup> | >64.0<sup>R</sup> |
| **PMB_4** | >128<sup>R</sup> | >128<sup>R</sup> | 4.00 | >64.0<sup>R</sup> | >64.0<sup>R</sup> | ≥8.00<sup>I,R</sup> | ≥32.0<sup>R</sup> | ≤2.00<sup>S</sup> | 48.00<sup>R</sup> | 2.00<sup>S</sup> | ≥8.00<sup>R</sup> | >64.0<sup>I</sup> | >64.0<sup>R</sup> | 2.00<sup>I</sup> | >64.0<sup>R</sup> |
| **PMB_5** | >128<sup>R</sup> | >128<sup>R</sup> | ≤8.00 | >64.0<sup>R</sup> | >64.0<sup>R</sup> | ≥8.00<sup>I,R</sup> | ≥32.0<sup>R</sup> | ≤8.00<sup>S</sup> | >64.0<sup>R</sup> | ≤4.00<sup>S</sup> | ≥2.00<sup>I,R</sup> | >64.0<sup>I</sup> | >64.0<sup>R</sup> | ≤2.00<sup>S,I</sup> | >64.0<sup>R</sup> |
| **PMB_6** | >128<sup>R</sup> | >128<sup>R</sup> | ≤8.00 | >64.0<sup>R</sup> | >64.0<sup>R</sup> | ≥16.0<sup>R</sup> | >64.0<sup>R</sup> | 8.00<sup>S</sup> | >64.0<sup>R</sup> | ≤4.00<sup>S</sup> | ≥32.0<sup>R</sup> | >64.0<sup>I</sup> | >64.0<sup>R</sup> | ≤4.00<sup>I,R</sup> | >64.0<sup>R</sup> |
| **OctC4_1** | 0.25<sup>S</sup> | ≤0.25<sup>S</sup> | 32.0 | >64.0<sup>R</sup> | >64.0<sup>R</sup> | ≥16.0<sup>R</sup> | >64.0<sup>R</sup> | 8.00<sup>S</sup> | >64.0<sup>R</sup> | ≤2.00<sup>S</sup> | ≥32.0<sup>R</sup> | >64.0<sup>I</sup> | >64.0<sup>R</sup> | ≤2.00<sup>S,I</sup> | >64.0<sup>R</sup> |
| **OctC4_2** | 0.25<sup>S</sup> | 0.25<sup>S</sup> | 32.0 | >64.0<sup>R</sup> | >64.0<sup>R</sup> | ≥8.00<sup>I,R</sup> | >64.0<sup>R</sup> | 8.00<sup>S</sup> | >64.0<sup>R</sup> | ≤2.00<sup>S</sup> | ≥32.0<sup>R</sup> | >64.0<sup>I</sup> | ≤2.00<sup>S</sup> | ≤1.00<sup>S</sup> | ≥8.00<sup>R</sup> |
| **OctC4_3** | ≤0.5<sup>S</sup> | 0.25<sup>S</sup> | 32.0 | >64.0<sup>R</sup> | >64.0<sup>R</sup> | ≤4.00<sup>S,I</sup> | 32.0<sup>R</sup> | 8.00<sup>S</sup> | >64.0<sup>R</sup> | ≤4.00<sup>S</sup> | ≤0.25<sup>S</sup> | >64.0<sup>I</sup> | >64.0<sup>R</sup> | 2.00<sup>I</sup> | >64.0<sup>R</sup> |
| **OctC4_4** | ≤0.5<sup>S</sup> | 0.25<sup>S</sup> | >32.0 | >64.0<sup>R</sup> | >64.0<sup>R</sup> | ≤4.00<sup>S,I</sup> | 32.0<sup>R</sup> | 8.00<sup>S</sup> | >64.0<sup>R</sup> | 1.00<sup>S</sup> | ≤0.25<sup>S</sup> | >64.0<sup>I</sup> | ≥2.00<sup>S,R</sup> | ≤2.00<sup>S,I</sup> | ≥4.00<sup>R</sup> |
| **OctC4_5** | 0.50<sup>S</sup> | 0.50<sup>S</sup> | 32.0 | >64.0<sup>R</sup> | >64.0<sup>R</sup> | >64.0<sup>R</sup> | >64.0<sup>R</sup> | ≤8.00<sup>S</sup> | >64.0<sup>R</sup> | 2.00<sup>S</sup> | >64.0<sup>R</sup> | >64.0<sup>I</sup> | >64.0<sup>R</sup> | 2.00<sup>I</sup> | >64.0<sup>R</sup> |
| **OctC4_6** | ≤0.5<sup>S</sup> | ≤0.5<sup>S</sup> | 32.0 | >64.0<sup>R</sup> | >64.0<sup>R</sup> | ≤4.00<sup>S,I</sup> | ≥32.0<sup>R</sup> | ≤16.0<sup>S,I</sup> | >64.0<sup>R</sup> | 2.00<sup>S</sup> | ≤0.25<sup>S</sup> | >64.0<sup>I</sup> | >64.0<sup>R</sup> | 2.00<sup>I</sup> | >64.0<sup>R</sup> |

<sup>a</sup>Initial polymyxin-susceptible isolate is 20\_GR\_12 compared against 20 days of treatment against polymyxin B (PMB), colistin (CST) or Octapeptin C4 (OctC4).
<sup>b</sup>Minimum Inhibitory Concentration tested for CST, Colistin; PMB, polymyxin B; OctC4, Octapeptin C4; AMX, Amoxicillin; ATM, Aztreonam; FEP, Cefepime; CRO, Ceftriaxone; CHL, Chloramphenicol; CIP, Ciprofloxacin; GEN, Gentamicin; MEM, Meropenem; PIP, Piperacillin; TET, Tetracycline; TGC, Tigecycline; TMP, Trimethoprim. Resistance determined as per CLSI guidelines and EUCAST for tigecycline with: S, Susceptible; I, Intermediate; R, Resistant. Fluctuations in MIC values (n=4) are displayed by two letters defining the resistance level. Resistance for Octapeptin C4 defined as $\geq 32$ $\mu\text{g/ml}$ .

### Octapeptin resistance induced isolates harbor an increase in hydroxymyristate and palmitoylate dissimilar to Ara4N lipid A modifications in Pmx-R strains

In the initial isolate, MS/MS analysis of extracted lipid A fractions showed that the major singly charged peak was *m/z* 1824.2, which corresponded to a hexa-acylated lipid A species comprised of two phosphate groups, two glucosamines and four 3-hydroxy-myristoyl groups (3-OH-C_14_), with two of these further acylated with myristate (C_14_) (Fig. 3A), also see Fig. S1 in the supplemental material). Due to the low intensity of this peak and the maximum detection limit of 2000 Da, in the system used, doubly charged masses were examined (see Fig. S1A in the supplemental material). This mass correlated to a doubly charged species of *m/z* 911.6 herein designated as the wild-type (WT) lipid A. Lesser quantities of various modifications accompanied the WT lipid A in the initial strain, including a hydroxyl modification of a myristate (*m/z* 919.6, WT+C_14:OH_), palmitoylation (*m/z* 1030.7 WT+C_16_) and even the addition of Ara4N (*m/z* 977.1, WT+Ara4N), a modification known to confer polymyxin resistance (Fig. 3B). In sharp contrast, the predominant species found in Pmx-R isolates was the near complete loss of WT lipid A and fortification of Ara4N on phosphate groups, mainly in hydroxymyristate species (*m/z* 985.1, WT+C_14:OH_+Ara4N; *m/z* 1042.7, WT+2(Ara4N); *m/z* 1050.7, WT+C_14:OH_;+(Ara4N)) (Fig. 3B), see Fig. S2 and S3 in the supplemental material). These changes corresponded to the genetic changes described in the following sections. The other commonly reported lipid A modification for resistance, pEtN, corresponding to *m/z* 973.2 was never observed. The lipid A from the OctC4 induced isolates was substantially different from the Pmx-R isolates and similar to the WT profile, with a major peak of the hydroxymyristate derivative and a significant 5-fold increase in representation of palmitoylation (Fig. 3B), see Fig. S4 in the supplemental material). The Ara4N modification was enhanced compared to WT, but not to the extent seen with Pmx-R isolates.

**FIG.3.**
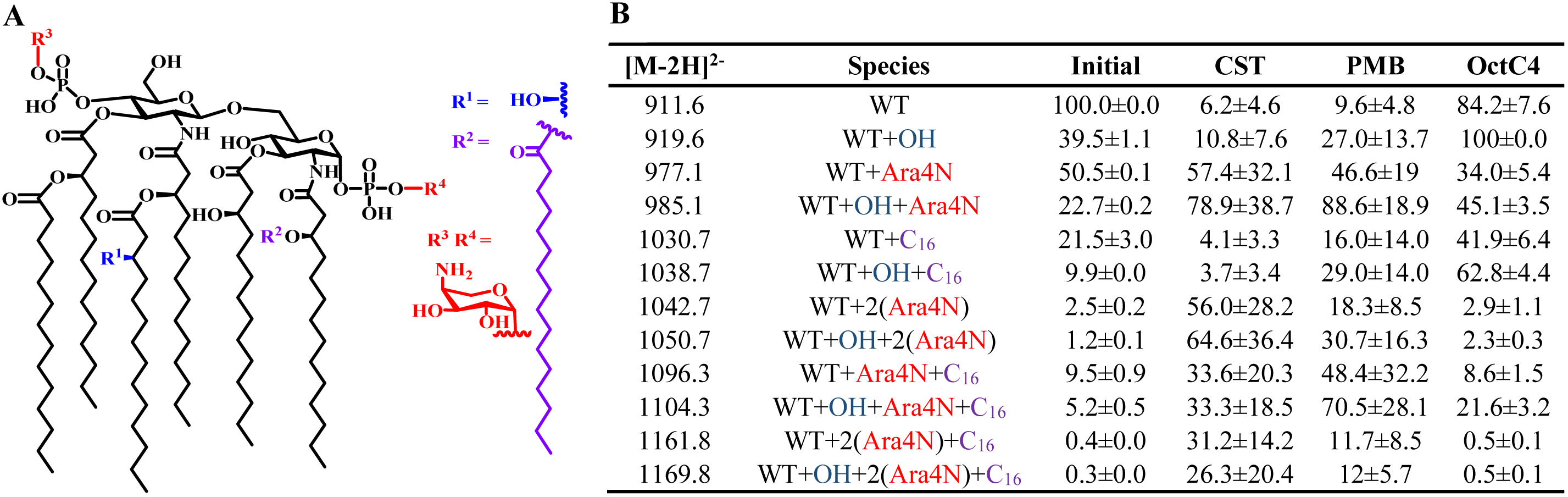
Lipid A modifications identified after 20 days of exposure to either colistin, polymyxin B or octapeptin C4. (A) Modifications which were detected in wild-type (WT) hexa-acylated lipid A. This included hydroxylation of a myristate (R^1^), palmitoylation (R^2^) and the addition of 4-amino-4-deoxy-arabinose (Ara4N) to either phosphate groups (R^3^, R^4^). (B) Doubly charged lipid A species detected for the initial isolate (n=2) and treatment groups (n=6). Values represent mean±SD of relative peak intensities.

### Plasmid loss associated with OctC4 resistance

To ascertain the genetic basis for resistance and subsequent phenotypic traits, four day 20 replicates were selected from each treatment group. Clonal expansion of genomic variations were monitored by selecting four colonies per replicate. Additionally, two colonies from the initial isolate were sequenced. The initial isolate harbored multiple acquired resistance genes targeting aminoglycosides, β-lactams, fosfomycin, quinolones, sulfonamides, tetracycline and trimethoprim, consistent with the parent XDR profile (Table 2). Five plasmid replicons were identified including ColRNAI, IncFIB(K)-Kpn3, IncFII(K), IncN and IncX3. In cross-resistance studies, the only Pmx-R replicate with an alteration in MIC profile to other antibiotics was PMB_2. Unique to PMB_2 was the susceptibility to meropenem and a reduction in resistance towards cefepime. Sequencing revealed a lack of *aph(3 ‘)-Ia, blaKPC-2, blaOXA-9* and no evidence of the IncX3 replicon in all four colonies. There was also a partial loss of this plasmid in PMB_4 (Table 2).

High variability of acquired resistance genes and plasmids were witnessed for OctC4 exposed replicates. Resistance genes impacted included *aph(3’)-Ia, aph(3”)-Ib, aph(6)-Id, blaKPC-2, blaOXA-9, blaTEM-1B, sul2, tet(A)* and *dfrA14*. Furthermore, plasmid replicon loss was apparent in three of the four replicates including IncFIB(K)-Kpn3, IncFII(K) and IncN. Subtle discrepancies in β-lactamase genes were observed across the three treatment groups; however, this was attributed to difficulties in the assembly due to high homology amongst these genes.

**TABLE 2.**
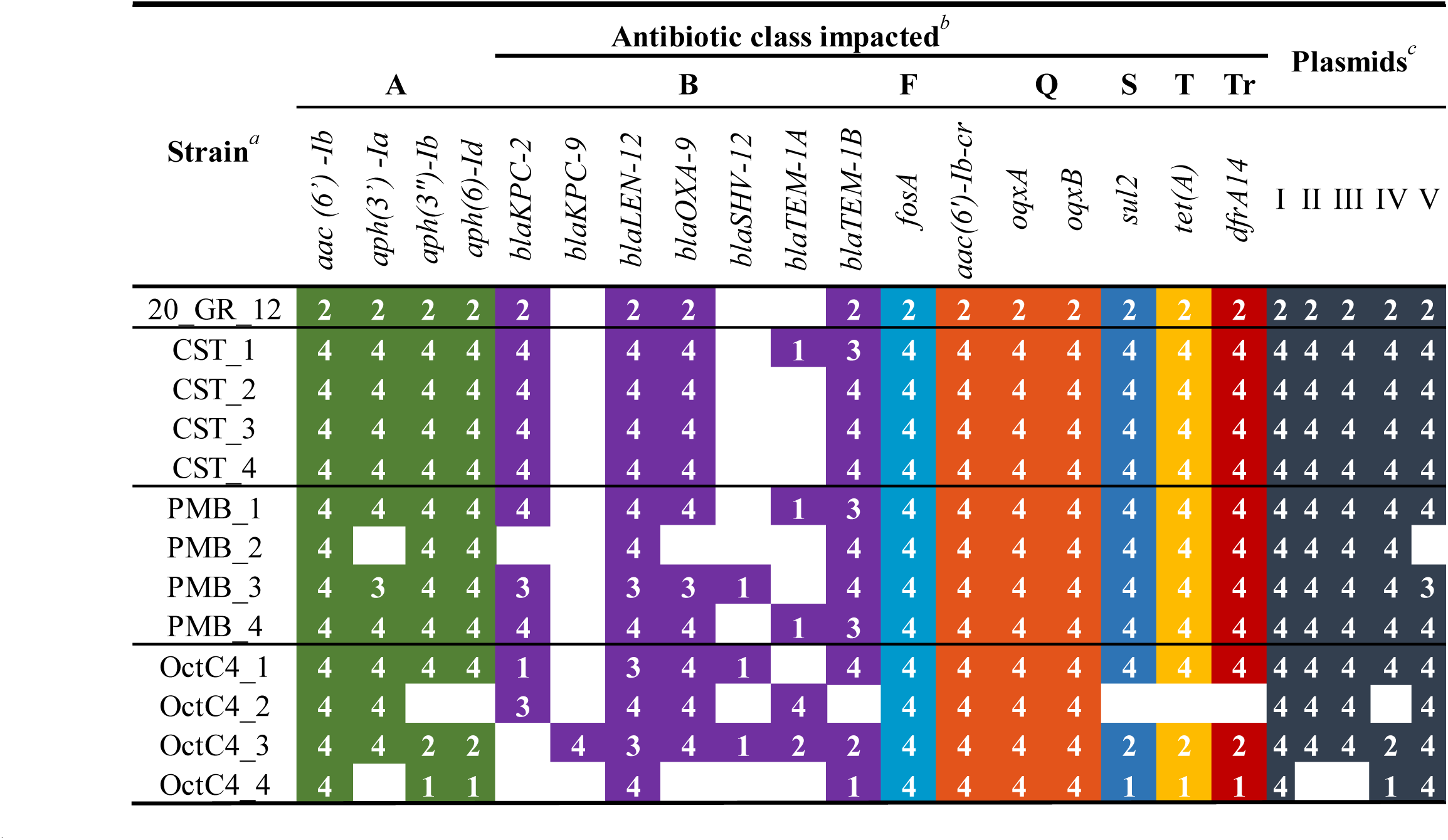
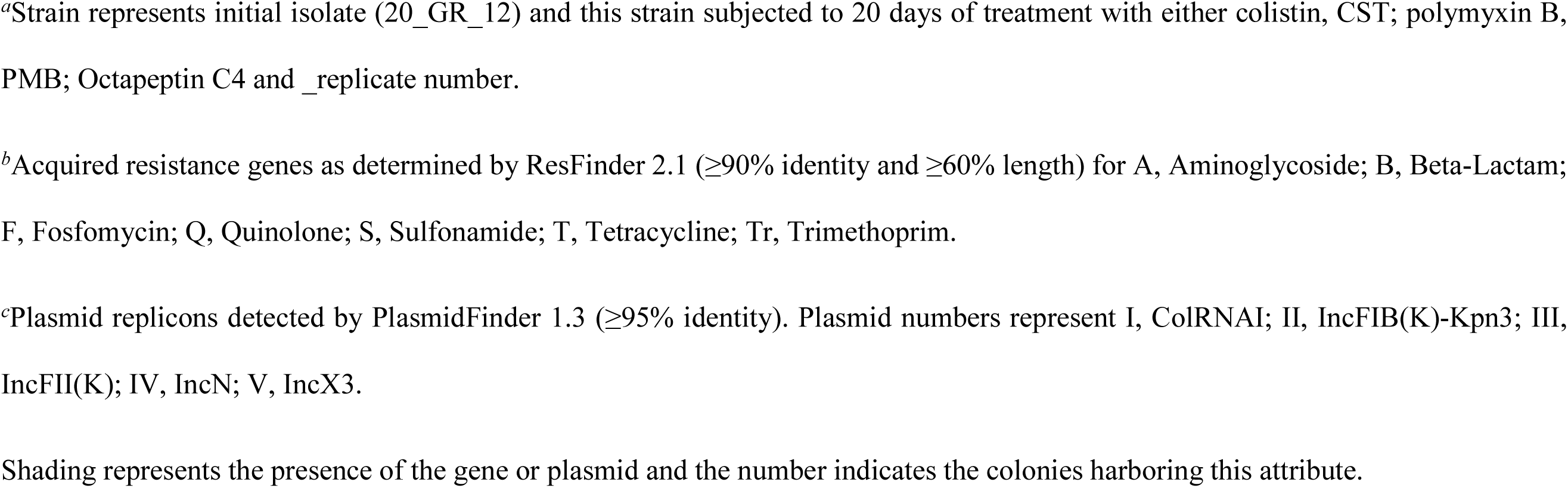
Detection of acquired resistance genes and plasmid replicons compared to the initial isolate and treatment groups

### Chromosomal variations in lipopolysaccharide pathways associated with polymyxin resistance whilst phospholipid transport associated with OctC4 resistance

Genomic alterations identified in polymyxin and OctC4 treated replicates differed significantly. In Pmx-R replicates, genes predominantly associated with LPS processing and lipid A modifications were altered, including *crrB, hepIII, lptC, mgrB, pmrB, phoPQ* and *yciM* (Table 3). An additional TCS gene, *qseC*, was also disrupted in PMB_3 (S8R, I283L) and PMB_4 (L40F). Although similar genes were impacted across replicates, the mutation positions differed. Additionally, an accumulation of variations in LPS pathways were apparent within a single replicate. All four colonies from a single replicate were commonly changed indicating clonal expansion of this variant. Complementation assays were further conducted to unveil the contributions of these genes to observed resistance (Fig. 4). Polymyxin susceptibility was restored in CST_2 (complete deletion of *mgrB)*, CST_3 (M1I), CST_4 (N42I), PMB_2 (W47L) and PMB_4 (D29Y) once complemented with pTOPO-mgrB (Fig. 4B-D, F and H). The PmrB (P95L) variant in PMB_1 was validated to contribute to resistance (Fig. 4E). Alterations in CrrB (D57V), PhoP (R81C) and QseC (S8R, I283L) were confirmed to cause resistance once these genes were introduced into the initial strain (Fig. 4M). Subtle increases in polymyxin MIC was detected for PhoQ (P420A, G434C), PhoQ (D417N) and QseC (L40F) but did not surpass the breakpoint (Fig. 4M). This confirms the presence of multiple resistance conferring mutations being present in a single isolate and several contributing to the elevation of MIC.

**FIG.4.**
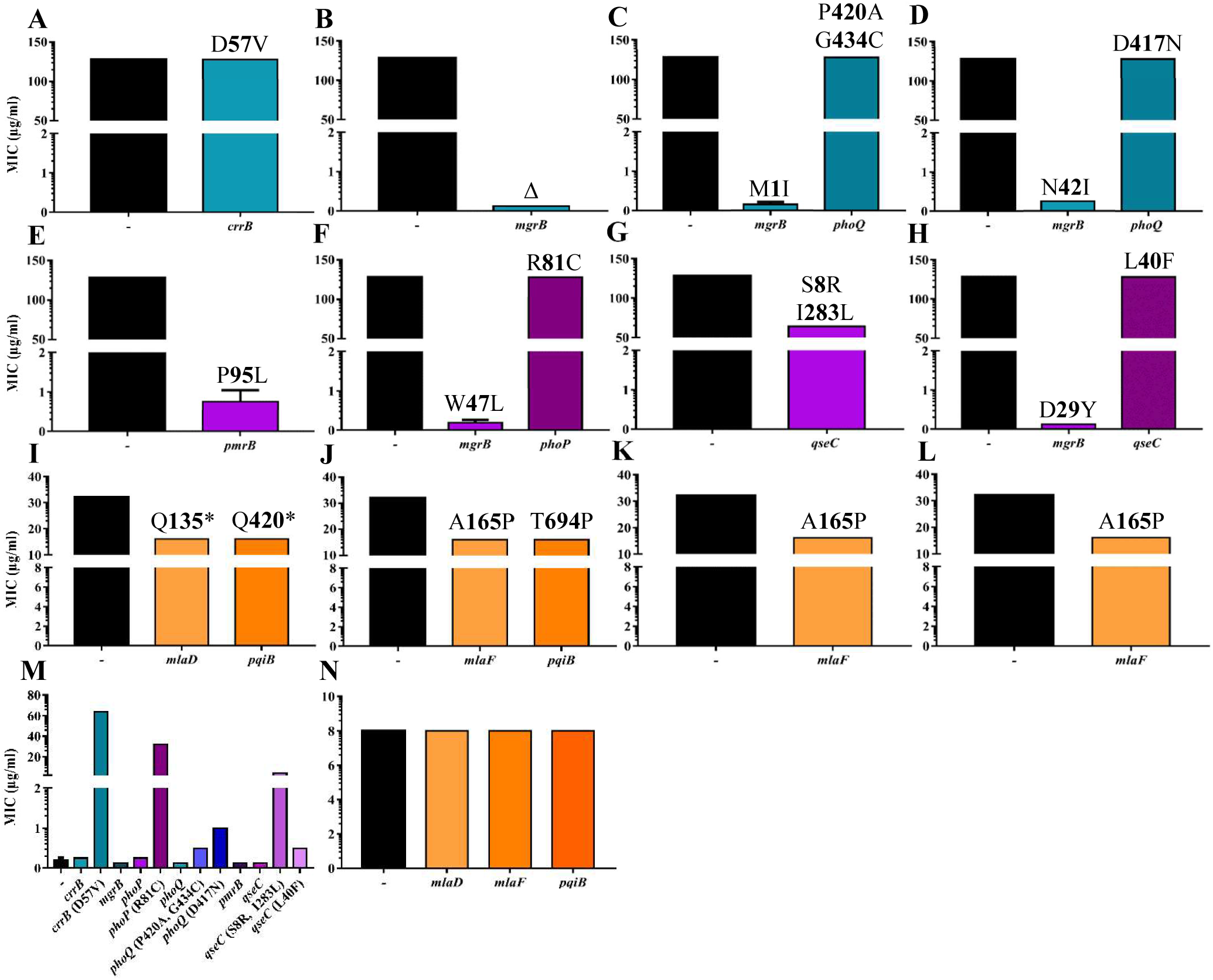
Complementation assays to delineate contribution of variations detected in day 20 treated strains to resistance. (A-D) Colistin treatment groups complemented with WT gene. (A) CST_1 with pTOPO-crrB. (B) CST_2 with pTOPO-mgrB. (C) CST_3 with pTOPO-mgrB or pTOPO-*phoQ*. (D) CST_4 with pTOPO-mgrB or *pTOPO-phoQ*. (E-H) Polymyxin B treatment groups complemented with WT gene. (E) PMB_1 with *pTOPO-pmrB*. (F) PMB_2 with pTOPO-mgrB or *pTOPO-phoP*. (G) PMB_3 with pTOPO-qseC. (H) PMB_4 with pTOPO-mgrB or pTOPO-qseC. (I-L) Octapeptin C4 treatment groups complemented with WT gene. (I) 0ctC4_1 with pTOPO-*mlaD* or *pTOPO-pqiB* (J) 0ctC4_2 with pTOPO-mlaF or *pTOPO-pqiB*. (K) 0ctC4_3 and with pTOPO-mlaF. (L) 0ctC4_4 with pTOPO-mlaF. (M) 20_GR_12, the initial strain, complemented with WT genes and genes harboring mutations potentially causing polymyxin resistance. (N) Complementation of octapeptin C4 resistance associated WT genes in 20_GR_12. The (-) indicates no complementation was conducted and represents the initial MIC. The y-axis split signifies the breakpoint for polymyxins (2 μg/ml) and initial MIC for octapeptin C4 (8 μg/ml). Values represented as mean±SD (n=4). Values above bars (A-L) indicate amino acid change in induced resistant isolate. Δ represents a complete deletion of protein and ^*^is a stop codon.

**TABLE 3.**
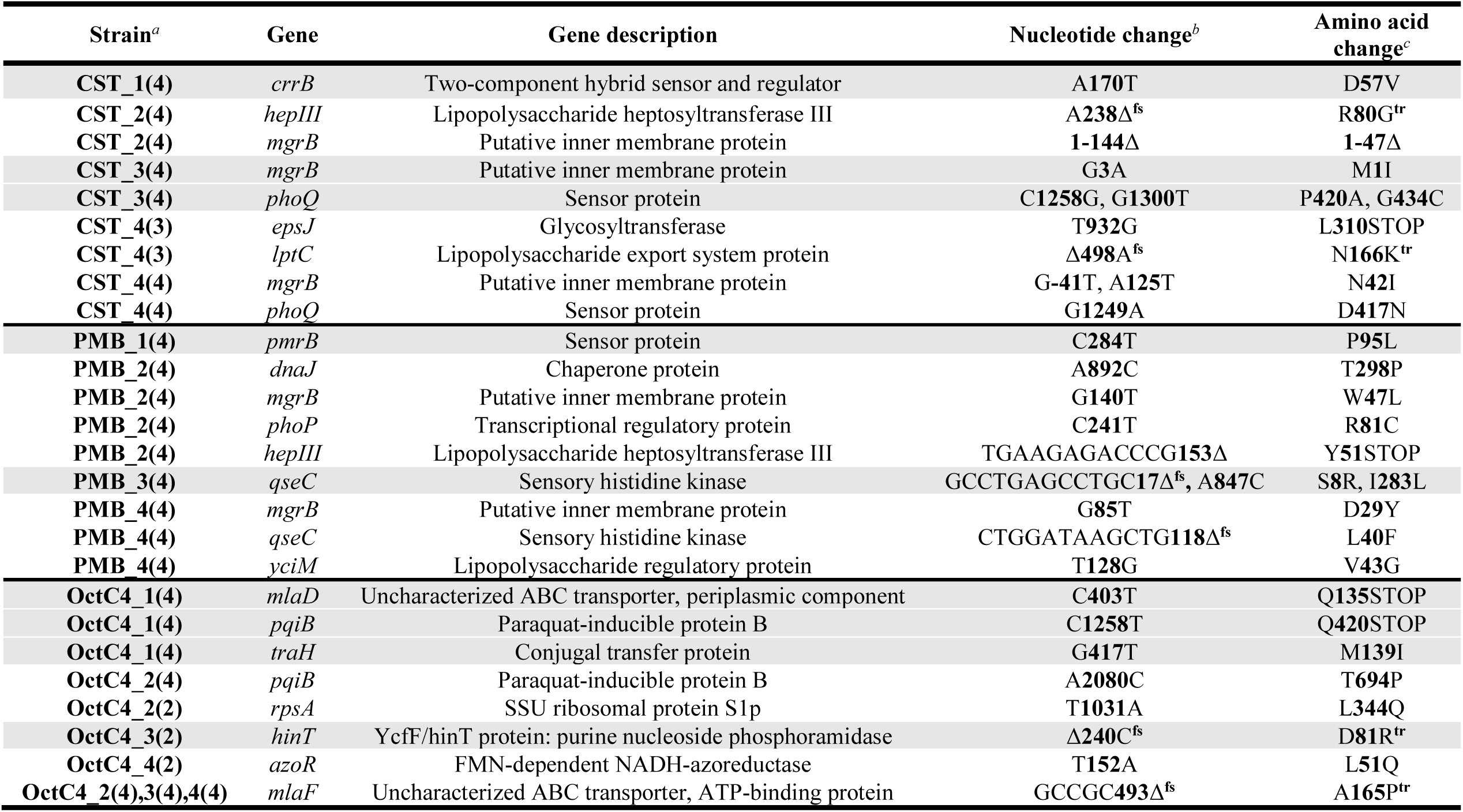
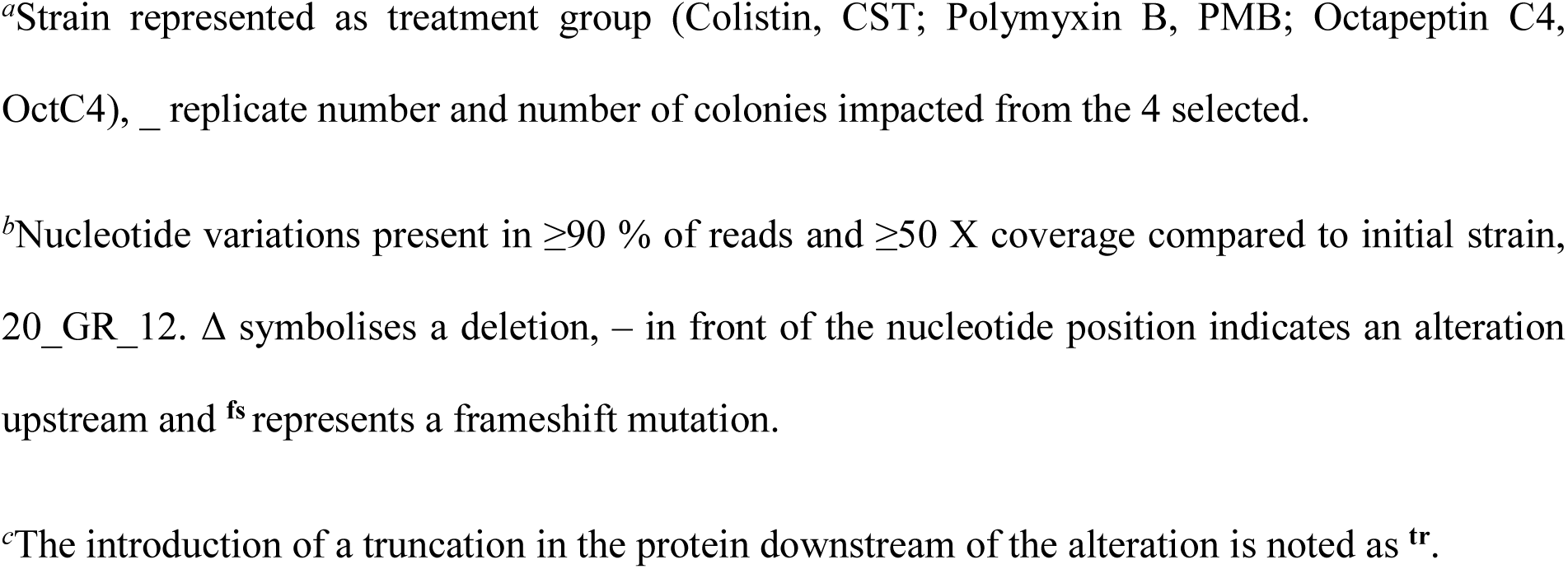
Genomic alterations detected in day 20 resistance induced isolates

| Strain<sup>a</sup> | Gene | Gene description | Nucleotide change<sup>b</sup> | Amino acid change<sup>c</sup> |
| --- | --- | --- | --- | --- |
| CST_1(4) | *crrB* | Two-component hybrid sensor and regulator | A170T | D57V |
| CST_2(4) | *hepIII* | Lipopolysaccharide heptosyltransferase III | A238Δ<sup>fs</sup> | R80G<sup>tr</sup> |
| CST_2(4) | *mgrB* | Putative inner membrane protein | 1-144Δ | 1-47Δ |
| CST_3(4) | *mgrB* | Putative inner membrane protein | G3A | M1I |
| CST_3(4) | *phoQ* | Sensor protein | C1258G, G1300T | P420A, G434C |
| CST_4(3) | *epsJ* | Glycosyltransferase | T932G | L310STOP |
| CST_4(3) | *lptC* | Lipopolysaccharide export system protein | Δ498A<sup>fs</sup> | N166K<sup>tr</sup> |
| CST_4(4) | *mgrB* | Putative inner membrane protein | G-41T, A125T | N42I |
| CST_4(4) | *phoQ* | Sensor protein | G1249A | D417N |
| PMB_1(4) | *pmrB* | Sensor protein | C284T | P95L |
| PMB_2(4) | *dnaJ* | Chaperone protein | A892C | T298P |
| PMB_2(4) | *mgrB* | Putative inner membrane protein | G140T | W47L |
| PMB_2(4) | *phoP* | Transcriptional regulatory protein | C241T | R81C |
| PMB_2(4) | *hepIII* | Lipopolysaccharide heptosyltransferase III | TGAAGAGACCCG153Δ | Y51STOP |
| PMB_3(4) | *qseC* | Sensory histidine kinase | GCCTGAGCCTGC17Δ<sup>fs</sup>, A847C | S8R, I283L |
| PMB_4(4) | *mgrB* | Putative inner membrane protein | G85T | D29Y |
| PMB_4(4) | *qseC* | Sensory histidine kinase | CTGGATAAGCTG118Δ<sup>fs</sup> | L40F |
| PMB_4(4) | *yciM* | Lipopolysaccharide regulatory protein | T128G | V43G |
| OctC4_1(4) | *mlaD* | Uncharacterized ABC transporter, periplasmic component | C403T | Q135STOP |
| OctC4_1(4) | *pqiB* | Paraquat-inducible protein B | C1258T | Q420STOP |
| OctC4_1(4) | *traH* | Conjugal transfer protein | G417T | M139I |
| OctC4_2(4) | *pqiB* | Paraquat-inducible protein B | A2080C | T694P |
| OctC4_2(2) | *rpsA* | SSU ribosomal protein S1p | T1031A | L344Q |
| OctC4_3(2) | *hinT* | YcfF/hinT protein: purine nucleoside phosphoramidase | Δ240C<sup>fs</sup> | D81R<sup>tr</sup> |
| OctC4_4(2) | *azoR* | FMN-dependent NADH-azoreductase | T152A | L51Q |
| OctC4_2(4),3(4),4(4) | *mlaF* | Uncharacterized ABC transporter, ATP-binding protein | GCCGC493Δ<sup>fs</sup> | A165P<sup>tr</sup> |

<sup>a</sup>Strain represented as treatment group (Colistin, CST; Polymyxin B, PMB; Octapeptin C4, OctC4), \_ replicate number and number of colonies impacted from the 4 selected.
<sup>b</sup>Nucleotide variations present in $\geq 90$ % of reads and $\geq 50$ X coverage compared to initial strain, 20\_GR\_12. $\Delta$ symbolises a deletion, – in front of the nucleotide position indicates an alteration upstream and <sup>fs</sup> represents a frameshift mutation.
<sup>c</sup>The introduction of a truncation in the protein downstream of the alteration is noted as <sup>tr</sup>.

The OctC4 replicates harbored changes in *mlaDF, pqiB* and *traH* in all four colonies. Additional genes altered that were apparent in two colonies per replicate included *azoR, hinT* and *rpsA*. Strikingly, *mlaF* (A165P) was impacted in three different OctC4 replicates at the same position (Table 3). Complementation assays that introduced pTOPO-mlaD, *-mlaF* or *-pqiB* into OctC4 induced replicates reduced the MIC by 2-fold, however, consistently only partial growth was observed at 8 μg/ml. This finding validates the partial contribution of these genes to resistance (Fig. 4I-L). Introduction of WT genes into the initial isolate revealed that the vector and gene did not influence the MIC and confirmed that these alterations are responsible for the resistance observed (Fig. 4M and N).

## DISCUSSION

Polymyxin unfortunately now induce high levels of resistance during therapeutic use, which is further compromised by suboptimal exposure in the clinic due to the risk of nephrotoxicity (29). Resistance in *K. pneumoniae* appears to be stable and incurs a minimal fitness cost (30, 31). These clinical characteristics were reflected in our study whereby once the isolate could tolerate 0.5 μg/ml of either CST or PMB, the clinical breakpoint was vastly exceeded within 48 h, well within the duration of clinical antibiotic therapy. This rapid induction of resistance was not observed for OctC4, in which only comparatively minor increases in MIC were observed. The slow progression in resistance profile could be an advantageous characteristic of OctC4 as a potential clinical intervention.

Following 20 days of increasing sub-lethal antibiotic exposure, no cross-resistance was apparent between polymyxins and OctC4. CST and PMB resulted in similar profiles with the only deviation seen in sample PMB_2 in which susceptibility to meropenem was regained. This is due to the absence of *blaKPC-2* and *blaOXA-9*. Additionally, the homogenous loss of the IncX3 plasmid was identified. Clinically, meropenem is being used in combination with polymyxins, and these results suggest that, in some cases, meropenem may overcome polymyxin resistance (32, 33). Furthermore, previous research has identified the loss of *blaKPC* plasmids in Pmx-R clinical isolates and suggests that this loss is due to a potential fitness cost (34). Our results show various accounts of plasmid loss in OctC4-exposed replicates, and this corresponded to a reduction in resistance towards cefepime, meropenem and tetracycline. Whether this resembles a fitness cost associated with OctC4 exposure or due to repeated passaging under no selective pressure for the genes harbored on these plasmids warrants further investigation.

Interestingly, resistance towards chloramphenicol was diminished in polymyxin and OctC4 exposed strains. Resistance towards chloramphenicol can arise from plasmid-encoded chloramphenicol acetyltransferases, alterations in the target 50S ribosomal subunit, or disruptions in porins and efflux pumps (35). The absence of acquired resistance genes and the lack of modifications in these regions may imply either a down-regulation of efflux pumps or an alternative resistance mechanism. The synergistic mechanism of polymyxins and chloramphenicol have been extensively studied; however, this finding potentially indicates a novel loss of chloramphenicol resistance upon gaining resistance towards these lipopeptides (36, 37), and is also seen with the octapeptins.

The mutations observed in polymyxin resistance induced ST258 strains can be compared to those we have previously identified in closely related polymyxin-resistant clinical ST258 isolates, 2_GR_12, 4_GR_12, 10_GR_13, 13_GR_14 and 14_GR_14 (24). As in this study, the vast majority of resistance was attributed to *mgrB* (60%), albeit not via an IS element disruption commonly observed in the clinic. Additional mutations were also identified in *phoPQ* accompanying the *mgrB* disruption, which was also apparent in this study (CST_3, CST_4, PMB_2). Other mutations in *crrB, mgrB, pmrB, phoPQ* and *yciM* in acquisition of polymyxin resistance have previously been described in resistant strains (16, 17, 38). Taken together, this indicates that resistance induction experiments have the capacity to induce genomic changes observed in the clinic.

These mutations lead to increased levels of Ara4N–modified lipid A, as observed in the lipid A analysis. Interestingly, the initial polymyxin-susceptible isolate exhibited Ara4N lipid A modifications which reveals a heteroresistant strain where a subpopulation of resistant bacteria exists within a phenotypically susceptible isolate. In several instances for polymyxin induced isolates, mutations were present in multiple genes within a replicate. CST_3 harbored a deleterious mutation in *mgrB* (M1I) and additional alterations in *phoQ* (P420A, G434C) increased tolerance to CST. This was also the circumstance for CST_4 (*mgrB:* N42I, *phoQ:* D417N). PMB_2 possessed a resistance conferring mutation in *mgrB* (W47L) and *phoP* (R81C). The notion that one alteration in TCS drives resistance, the circumstance for the majority of clinical isolates is well accepted (39). However, our findings contradict this concept.

We also identified alterations in another TCS, QseBC, which is known to facilitate cross-talk with PmrAB in *Escherichia coli* (40). In *E. coli*, PmrB acts as a noncognate partner to the QseBC TCS and has the capability to not only phosphorylate PmrA, but also QseB. The absence of QseC was shown to impact virulence due to the accumulation of phosphorylated QseB and in particular, alterations in the histidine kinase domain attenuates its ability to de-phosphorylate QseB (40, 41). Furthermore, the deletion of *qseC* and *pmrA*, promoting phosphorylation of QseB by PmrB, stimulated tolerance to PMB (42). This signalling pathway remains severely under characterized in *K. pneumoniae*. We observed partial tolerance to PMB when a frameshift mutation was apparent at nucleotide 118; however, full resistance in PMB_4 was promoted by alterations in *mgrB* (D29Y) and *yciM* (V43G), which has recently been identified to cause resistance (38). Conversely, PMB_3 also harbored a frameshift mutation early in the coding sequence of *qseC* (GCCTGAGCCTGC17Δ^fs^), although an additional I283L change in the histidine kinase region resulted in an MIC of 4 μg/ml. This did not explain the full resistance profile exhibited by PMB_3 and due to the presence of both alleles during complementation, the true extent of resistance cannot be deduced. Considering PMB_3 still resulted in the addition of Ara4N to lipid A, we speculate that due to the perturbation in the QseC kinase, this is increasing the accumulation of phosphorylated QseB and allows for the up-regulation of transcriptional targets. Subsequent transcription could be activating PmrA, similar to other TCS in *K. pneumoniae*, allowing for the expression of the *pmrHFIJKLM* operon (Fig. 5A).

**FIG.5.**
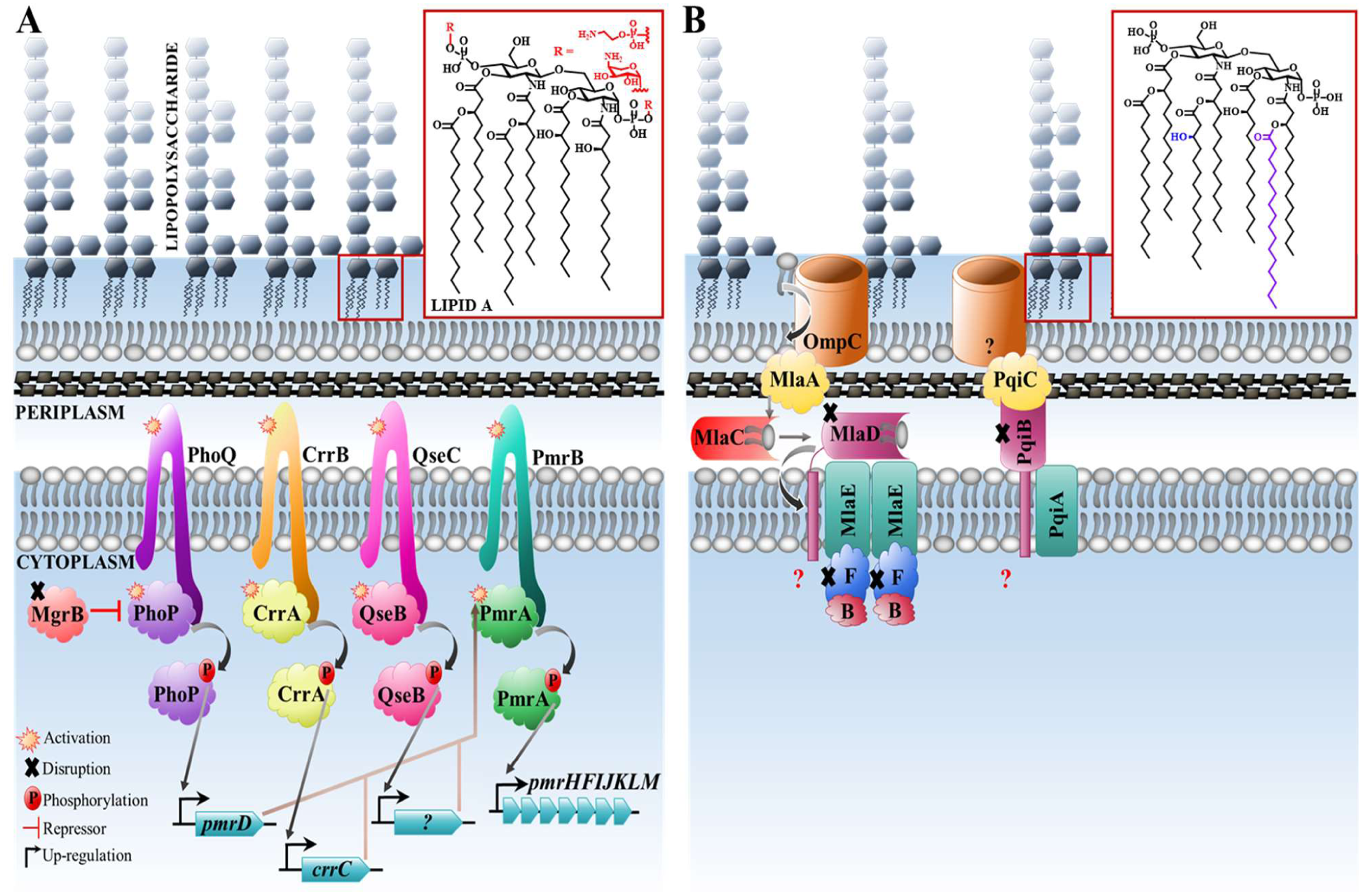
Proposed pathway associated with *K. pneumoniae* polymyxin and octapeptin C4 resistance observed in this study. (A) To facilitate resistance against polymyxins, genomic variations are acquired in two-component regulatory systems. These encompass CrrAB, QseBC, PmrAB and PhoPQ with MgrB acting as a negative repressor. Once this pathway is activated during resistance, sensor histidine kinases (CrrB, QseC, PmrB, PhoQ) will phosphorylate response regulators (CrrA, QseB, PmrA, PhoP) and allow for the expression of target genes (crrC, unknown (?), *pmrD, pmrHFIJKLM)*. Disruptions in MgrB allow for the up-regulation of this pathway resulting in the expression of *pmrHFIJKLM* which allows for 4-amino-4-deoxy-arabinose or phosphoethanolamine to be attached to phosphate groups on lipid A. (B) The major disruptions identified during octapeptin C4 resistance was in the Mla and Pqi pathway. OmpC removes phospholipids (PLs) from the outer member and transfers these to MlaA. PLs are transported across the periplasm via MlaC and transported to the MlaBDEF complex where the subsequent fate of PLs is unknown. An unknown porin complexes with PqiC to transport metabolites and potentially PLs across the periplasm via the PqiAB complex. Perturbations in these pathways elevated the MIC towards octapeptin C4 and subsequent hydroxymyristae and palmitoylation of lipid A to potentially stabilise the outer membrane.

The mutation pattern was greatly different in OctC4-exposed replicates, with all harboring alterations in the Mla pathway. These genes are responsible for phospholipid (PL) importation from the outer membrane into the cell (43). Removal of *mlaC* in *E. coli* was previously identified to increase the abundance of palmitoylated lipid A to stabilise the outer membrane which correlated to the phenotype in our study. Similarly, prior research exposing *P. aeruginosa* to OctC4 (32 μg/ml) revealed an increase in palmitoylated lipid A (22). Literature reports have demonstrated that octapeptins have the capacity to bind to PLs (44), and likely OctC4 utilises this pathway in order to traverse to the outer membrane (Fig. 5B). The involvement of PqiB in membrane integrity has only recently been characterized in *E. coli* (45). PqiB was identified to connect to PqiC and potentially deliver substrate/s from the outer to inner membrane. The contribution of the PqiABC appeared minimal compared to the Mla pathway and was proposed to either ineffectively transport PLs or transport different substrates with a minute impact on membrane integrity. The contribution of the Pqi and Mla pathway appeared to be additive when evaluating the MIC reduction in OctC4_1 and OctC4_2. Further genes impacted not homogeneous amongst the colonies included *rpsA* (40S ribosomal protein), *azoR* (quinone reductase), *traH* (plasmid conjugal transfer protein) and *hinT* (purine nucleoside phosphoramidase), which may indicate several intracellular targets (46-49). The lack of mutations associated with Ara4N-modifications to lipid A is consistent with the lipid A profile of the OctC4-induced isolates. This observation supports the hypothesis that the octapeptins work by a different mode of action compared to the polymyxins, one that does not require an initial binding to lipid A and explains the lack of cross-resistance between the two classes of lipopeptides. However, further studies are required to determine if this occurs ubiquitously for *K. pneumoniae* and if the same phenomenon is observed for other Gram-negative pathogens. The slow progression of resistance, potential fitness cost if resistance develops, and the alternative mechanism of infiltration of OctC4 highlight the potential for octapeptins to be explored as future antibiotics.

## MATERIALS AND METHODS

### Bacterial strains and growth conditions

Clinical polymyxin-susceptible XDR *K. pneumoniae* ST258 isolate, 20_GR_12, was sourced through Hygeia General Hospital, Athens, Greece as previously described (24). Cultures were grown in Luria-Bertani (LB) broth and for single colony isolation, cultures were grown on either LB or Nutrient Agar (NA) plates.

### Antimicrobial susceptibility assays

Minimum inhibitory concentration was determined by the broth microdilution method according to Clinical & Laboratory Standards Institute (CLSI) guidelines (50). Cultures were grown in cation-adjusted MHB and to assess cross-resistance of day 20 isolates, broth was supplemented with the concentration of antibiotic tolerated at that time point (see Table S1 in the supplemental material). Clinical breakpoints were determined in concordance to CLSI guidelines (51) and tigecycline as per The European Committee on Antimicrobial Susceptibility Testing (EUCAST) (Version 8.0, 2018) (see http://www.eucast.org).

### Induction of resistance

A single colony of the clinical isolate was selected and grown overnight at 37°C shaking at 220 rpm. Similar to the broth microdilution assay, this culture was grown to log phase (OD_600_ = 0.4-0.6). The culture was plated out into three separate 96-well polystyrene, non-treated plates (Sigma Aldrich) with six replicates for each treatment group including CST, PMB and OctC4. Plates were incubated overnight and OD_600_ was read at 20 h. The well which harbored dense growth (OD_600_ = %1) underwent a 1:1000 dilution, transferred to a new plate with the concentration range adjusted accordingly. The highest concentration used for the polymyxins was 128 μg/ml and 32 μg/ml for OctC4. This process was completed for 20 days with five following days of no antibiotic exposure. At day 20, the culture was further diluted (1:1000) and placed in non-supplemented broth to be incubated overnight. Part of this culture underwent an MIC against the antibiotic it was exposed to in order to evaluate stability of resistance. Several time points were isolated and stored in 30% sterile glycerol at -80°C for future assays. Fold change significance was determined via GraphPad Prism 7 with a one-way ANOVA with a Tukey’s multiple comparisons test where significance was P<0.05.

### Lipid A modifications

Lipid A was extracted using the ammonium hydroxide-isobutyric acid protocol as previously described (52). Day 20 cultures were grown overnight in LB supplemented with antibiotic (see Table S1 in the supplemental material). Overnight inoculums were subcultured (1:100) into 100 mL LB broth and grown to an OD_600_ = 0.8-1. Cultures were pelleted (10,000 rpm, 20 min, 4°C), washed with 1X PBS (10,000 rpm, 15 min, 4°C) and freeze dried. 10 mg of lyophilised cells were processed as per (52). Concisely, samples were suspended in isobutyric acid:ammonium hydroxide (5:3 [vol/vol]), under magnetic stirring at 100°C for 4 h, supernatants isolated by centrifugation at 13,000 rpm for 15 min, diluted with equal volume of water and freeze dried. Extracts then underwent two methanol washes (4,000 rpm, 15 min). The extracted lipid A was solubilised in methanol containing 5 mM ammonium acetate to a concentration of 1 μg/ml. Samples were infused at a low rate of 5 μl/min into a QSTAR Elite (Applied Biosystems) hybrid quadrupole Time-of-Flight (TOF) mass spectrometer. To acquire adequate fragmentation for MS/MS analysis, the collision energy was increased from 40 to 80. Averaged spectra were accumulated over at least 1 min. Data were exported from Analyst (SCIEX), normalized to the highest mass intensity and graphed in GraphPad Prism 7.

### DNA extractions and library preparation

Glycerol stocks from day 20 isolates were grown on NA plates overnight. Single colonies were isolated, grown in antibiotic supplemented broth (see Table S1 in the supplemental material), incubated overnight and DNA extracted using the DNeasy Blood and Tissue Kit (Qiagen) according to manufacturer’s guidelines. Two colonies were selected from day 0 and 4 colonies from 4 replicates per treatment group. Quantification of DNA was acquired using Qubit®3.0 (ThermoFisher Scientific) and 1 ng of DNA underwent library preparation with the Nextera XT kit (Illumina) as per manufacturer’s instructions. Quality control was checked with a 2100 Bioanalyzer (Agilent Technologies) and LabChip GX (PerkinElmer).

### Sequencing and analysis

Libraries were sequenced on an Illumina NextSeq with 150 bp paired end sequencing reads with ≥95X coverage with the exception of CST_2 (colony 1) (48X). Trimmomatic (53) was used to trim paired end reads and SPAdes v3.10.1 implemented for assembly (54). Annotation of assembled genomes was accomplished using the Rapid Annotation using Subsystem Technology (RAST) (55). The Centre for Genomic Epidemiology (CGE) tools were implemented to delineate laterally acquired resistant genes (ResFinder 3.0) (56) and plasmids (PlasmidFinder 1.3) (57). Reads were aligned using BWA-MEM (58), analyzed through FreeBayes (59) and impact of change determined through snpEff (60). Nucleotide sequences have been deposited under NCBI BioProject PRJNA415530 (www.ncbi.nlm.nih.gov/bioproject/415530).

### Complementation assays

Genes speculated to cause resistance underwent complementation as previously described (61). Briefly, genes harboring a potential variation contributing to resistance was amplified using the 2X Phusion HF master mix (ThermoFisher) with primers listed in Table S2 in the supplemental material. The gene was cloned into the pCR-BluntII-TOPO using the Zero Blunt TOPO PCR cloning kit (Invitrogen). The plasmid was transformed in electrocompetent *E. coli* T0P10 via electroporation, grown overnight on MHB agar supplemented with kanamycin (50 μg/ml) at 37°C overnight and plasmids extracted using the QIAprep spin miniprep column kit (Qiagen). Plasmids were transformed into the initial susceptible strain (20_GR_12) and incubated overnight on MHB agar containing zeocin (1500 μg/ml). Furthermore, the wild-type gene was amplified from the initial strain and placed into the resistant day 20 isolates. An MIC was conducted to determine if resistance was altered as mentioned above.

## ACKNOWLEDGEMENTS

MAC is an NHMRC Principal Research Fellow (APP1059354) and currently holds a fractional Professorial Research Fellow appointment at the University of Queensland with his remaining time as CEO of Inflazome Ltd. a company headquartered in Dublin, Ireland that is developing drugs to address clinical unmet needs in inflammatory disease by targeting the inflammasome. LJMC is an ARC Future Fellow (FT110100972). MEP is an Australian Postgraduate Award scholar. MATB is supported in part by a Wellcome Trust Strategic Award 104797/Z/14/Z. Research was supported by NHMRC grants APP1005350, APP1045326 and NIH grant R21AI098731/R33AI098731-03. We would like to acknowledge Dr Ilias Karaiskos and Dr Helen Giamarellou for supplying this clinical isolate and Dr Alysha Elliott for the initial characterization of the strain. We thank Dr Alejandra Gallardo-Godoy, Dr Karl Hansford, David Edwards and Ruby Pelingon for the synthesis and purification of OctC4, Dr Tim Bruxner and Angelika Christ for their support with the sequencing, Alun Jones with his assistance for mass spectrometry which was acquired at the IMB Mass Spectrometry Facility at UQ, and Dr Hannah Sidjabat for her assistance with the initial quality control of the resistance induced strains.

## Author contributions

MEP, MAC, LJMC, MATB conceived this study. MEP, MDC and DG performed the sequencing analysis and MEP, MSB and SR performed the experiments. MEP wrote the paper with the input from other authors.

